# Sensitivity and Specificity of Information Criteria

**DOI:** 10.1101/449751

**Authors:** John J. Dziak, Donna L. Coffman, Stephanie T. Lanza, Runze Li, Lars S. Jermiin

## Abstract

Information criteria (ICs) based on penalized likelihood, such as Akaike’s Information Criterion (AIC), the Bayesian Information Criterion (BIC), and sample-size-adjusted versions of them, are widely used for model selection in health and biological research. However, different criteria sometimes support different models, leading to discussions about which is the most trustworthy. Some researchers and fields of study habitually use one or the other, often without a clearly stated justification. They may not realize that the criteria may disagree. Others try to compare models using multiple criteria but encounter ambiguity when different criteria lead to substantively different answers, leading to questions about which criterion is best. In this paper we present an alternative perspective on these criteria that can help in interpreting their practical implications. Specifically, in some cases the comparison of two models using ICs can be viewed as equivalent to a likelihood ratio test, with the different criteria representing different alpha levels and BIC being a more conservative test than AIC. This perspective may lead to insights about how to interpret the ICs in more complex situations. For example, AIC or BIC could be preferable, depending on the relative importance one assigns to sensitivity versus specificity. Understanding the differences and similarities among the ICs can make it easier to compare their results and to use them to make informed decisions.

**Key Points:**

- Information criteria such as AIC and BIC are motivated by different theoretical frameworks.
- However, when comparing pairs of nested models, they reduce algebraically to likelihood ratio tests with differing alpha levels.
- This perspective makes it easier to understand their different emphases on sensitivity versus specificity, and why BIC but not AIC possesses model selection consistency.
- This perspective is useful for comparisons, but it does not mean that the information criteria are only likelihood ratio tests. Information criteria can be used in ways these tests themselves are not as well suited for, such as for model averaging.

## 1 Introduction

Many model selection techniques have been proposed for many different settings (see Claeskens and Hjort, 2008). Among other considerations, researchers must balance sensitivity (suggesting enough parameters to accurately model the patterns, processes, or relationships in the data) with specificity (not suggesting nonexistent patterns, processes, or relationships). Several of the simplest and most popular model selection criteria can be discussed in a unified way as log-likelihood functions with simple penalties. These include Akaike’s Information Criterion (Akaike, 1973, AIC), the Bayesian Information Criterion (Schwarz, 1978, BIC), the sample-size-adjusted AIC or AIC_c_ of Hurvich and Tsai (1989), the “consistent AIC” (CAIC) of Bozdogan (1987), and the sample-size-adjusted BIC (ABIC) of Sclove (1987) (see Table 1). Each of these ICs consists of a goodness-of-fit term plus a penalty to reduce the risk of overfitting, and each provides a standardized way to balance sensitivity and specificity. These criteria are widely used in model selection in many different areas, such as choosing network models for gene expression data in molecular phylogenetics (Darriba *et al.*, 2012; Edwards *et al.*, 2010; Jayaswal *et al.*, 2014; Kalyaanamoorthy *et al.*, 2017; Lefort et *al.*, 2017; Luo *et al.*, 2010; Posada and Buckley, 2004; Posada, 2008, 2009), in selecting covariates for regression equations (Miller, 2002), and in choosing the number of subpopulations in mixture models (Nylund *et al.*, 2007). In addition to being used as measures of fit for directly comparing models, they are also used as ways of tuning or weighting more complicated and specialized methods (e.g. Minin *et al.*, 2003; Bouveyron and Brunet-Saumard, 2014) such as automated model search algorithms in high-dimensional modeling settings where comparison of each possible model separately might be too difficult (e.g. Wang *et al.*, 2007a). For these reasons, it is widely useful to understand their rationale and relative performance.

**Table 1:** Summary of Common Information Criteria

| Criterion | Penalty Weight | Emphasis | Likely Kind of Error |
| --- | --- | --- | --- |
| Non-consistent criteria |  |  |  |
| AIC | $A_n = 2$ | Good prediction | Overfitting |
| AIC<sub>c</sub> | $A_n = 2n/(n - p - 1)$ | Good prediction | Overfitting |
| Consistent criteria |  |  |  |
| ABIC | $A_n = \ln((n + 2)/24)$ | Depends on $n$ | Depends on $n$ |
| BIC | $A_n = \ln(n)$ | Parsimony | Underfitting |
| CAIC | $A_n = \ln(n) + 1$ | Parsimony | Underfitting |

**Notes.** AIC = Akaike information criterion. ABIC = adjusted Bayesian information criterion. BIC = Bayesian information criterion. CAIC = Consistent Akaike information criterion. $n$ = sample size (number of subjects). Other criteria include the DIC (deviance information criterion) which acts as an analog of AIC in certain Bayesian analyses but is more complicated to compute.

Model selection using an IC involves choosing the model with the best penalized log-likelihood: that is, the highest value of *ℓ – A_n_p*, where *ℓ* is the log-likelihood of the entire dataset under the model, where *A_n_* is a constant or a function of the sample size *n*, and where *p* is the number of parameters in the model. For historical reasons, instead of finding the highest value of *ℓ* minus a penalty, this is often expressed as finding the lowest value of *–2ℓ* plus a penalty:

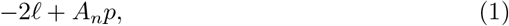

and we follow that convention here. This function is often computed automatically by computer software. However, to avoid confusion, investigators should be careful when using statistical software to be sure of what form is being used; in this paper we use the form in which the smaller IC is better, but if *ℓ – A_n_p* is used then the larger IC is better. Also, the form of the likelihood function and the definition of the parameters depends on the nature of the model. For example, in linear regression, *ℓ* is the multivariate normal log-likelihood of the sample, and *–2ℓ* becomes equivalent to *n* log(MSE) plus a constant, where MSE is the mean of squared prediction errors; *p* in this context is the number of regression coefficients. In latent class models, the likelihood is given by a multinomial distribution, and the parameters may include the means of each class on each dimension of interest and the sizes of the classes.

Expression (1) is what Atkinson (1980) called the generalized information criterion; in this paper we simply refer to (1) as an IC. Expression (1) is sometimes replaced in practice by the practically equivalent *G*^2^ + *A_n_p*, where *G*^2^ is the deviance, defined as twice the difference in log-likelihood between the current model and the saturated model, that is, the model with the most parameters which is still identifiable (e.g., Collins and Lanza, 2010).

In practice, Expression (1) cannot be used directly without first choosing *A_n_*. Specific choices of *A_n_* make (1) equivalent to AIC, BIC, ABIC or CAIC. Thus, although motivated by different theories and goals, algebraically these criteria are only different values of *A_n_* in (1), corresponding to different relative degrees of emphasis on parsimony, that is, on the number of free parameters in the selected model (Claeskens and Hjort, 2008; Lin and Dayton, 1997; Vrieze, 2012). Because the different ICs often do not agree, the question often arises as to which is best to use in practice.

For example, Miaskowski *et al.* (2015) recently used a latent class approach to categorize cancer patients into empirically defined clusters based on the presence or absence of 13 self-reported physical and psychological symptoms. They then showed that these clusters differed in terms of other covariates and on quality of life ratings, and suggested that they might have different treatment implications. Using BIC, they determined that a model with 4 classes (low physical symptoms and low psychological symptoms; moderate physical and low psychological; moderate physical and high psychological; high physical and high psychological) fit the data best. Their use of BIC was a very common choice and was recommended by work such as Nylund *et al.* (2007). It was not an incorrect choice, and we emphasize that we are not arguing that their results were flawed in any way. However, the AIC, ABIC, and CAIC, can be calculated from the information they provide in their Table 1, and if they had used AIC or ABIC it appears that they would have chosen at least a 5-class model instead. On the other hand, CAIC would have agreed with BIC. Does this mean that two of the criteria are incorrect and two are correct? We argue that neither is wrong, even though in their case the authors had to choose one or the other.

For a similar example using familiar and easily accessed data, consider the famous “Fisher’s iris data,” a collection of four measurements (sepal length, sepal width, petal length, petal width) of 50 flowers from each of 3 species of iris *(Iris setosa, Iris versicolor*, and *Iris virginica*). This data, originally collected by Anderson (1935) and famously used in an example by the influential statistician R. A. Fisher, is available in the dataset package as part of the base installation in R (R Core Team, 2017). It is often used for benchmarking and comparing statistical methods. For example, one can try clustering methods to classify the 150 flowers into latent classes without reference to the original species label and using only their measurements, and determine whether the methods correctly separate the three species. For a straightforward estimation approach (Gaussian model-based clustering without assuming equal covariance matrices; code is shown in the appendix), AIC or ABIC choose a 3-class model and BIC or CAIC choose a 2-class model. In the 3-class model, each of the empirically estimated classes corresponds almost perfectly to one of the three species, with very few misclassifications (5 of the *versicolor* were mistakenly classified as *virginica*). In the 2-class model, the *versicolor* and *virginica* flowers were lumped together. Agusta and Dowe (2003) performed this analysis and concluded that BIC performed poorly on this benchmark dataset. Most biologists would probably agree with this assessment. However, an alternative interpretation might be that BIC was simply being parsimonious, and that flower dimensions alone might not be enough to confidently separate the species. A much more detailed look at clustering the iris data, considering many more possible modeling choices, is found in Kim and Seo (2014). However, this simple look is enough to discuss the relevant ideas.

In this review we examine the question of choosing a criterion by focusing on the similarities and differences among AIC, BIC, CAIC, and ABIC, especially in view of an analogy between their different complexity penalty weights *A_n_* and the α levels of hypothesis tests. We especially focus on AIC and BIC, which have been extensively studied theoretically (Ding *et al.*, 2018; Kadane and Lazar, 2004; Kuha, 2004; Shao, 1997; Vrieze, 2012), and which are not only often reported directly as model fit criteria, but also used in tuning or weighting to improve the performance of more complex model selection techniques (e.g., in high-dimensional regression variable selection; Narisetty and He, 2014; Wang *et al.*, 2007b; Wu and Ma, 2015).

In the following section we review the motivation and theoretical properties of these ICs. We then discuss their application to a common application of model selection in medical, health and social scientific applications: that of choosing the number of classes in a finite mixture (latent class) analysis (e.g., Collins and Lanza, 2010). Finally, we propose practical recommendations for using ICs to extract valuable insights from data while acknowledging their differing emphases.

### 1.1 Common Penalized-Likelihood Information Criteria

In this section we review some commonly used ICs. Their formulas, as well as some of their properties which we describe later in the paper, are summarized for convenience in Table 1.

#### 1.1.1 Akaike’s Information Criterion (AIC)

First, the AIC Akaike (1973) sets *A_n_* = 2 in (1). It estimates the relative Kullback-Leibler (KL) divergence (a nonparametric measure of difference between distributions) of the likelihood function specified by a fitted candidate model, from the likelihood function governing the unknown true process that generated the data. The fitted model closest to the truth in the KL sense would not necessarily be the model that best fits the observed sample, since the *observed* sample can often be fit arbitrary well by making the model more and more complex. Rather, the best KL model is the model that most accurately describes the population distribution or the process that produced the data. Such a model would not necessarily have the lowest error in fitting the data already observed (also known as the training sample) but would be expected to have the lowest error in predicting future data taken from the same population or process (also known as the test sample). This is an example of a bias-variance tradeoff (see, e.g., Hastie *et al.*, 2009).

Technically, the KL divergence can be written as *E_t_(ℓ_t_(y)) – E_t_(ℓ(y))*, where *E_t_* is the expected value under the unknown true distribution function, *ℓ* is the log-likelihood of the data under the fitted model being considered, and *ℓ_t_* is the log-likelihood of the data under the unknown true distribution. This is intuitively understood as the difference between the estimated and the true distribution. *E_t_ (ℓt* (*y*)) will be the same for all models being considered, so KL is minimized by choosing the model with highest *E_t_(ℓ(y)*). The *ℓ(y)* from the fitted model is a biased measure of *E_t_(ℓ(y)*), especially if *p* is large, because a model with many parameters can generally be fine-tuned to appear to fit a small dataset well, even if its structure is such that it cannot generalize to describe the process that generated the data. Intuitively, this means that if there are many parameters, the fit of the model to the originally obtained data (training sample) will seem good regardless of whether the model is correct or not, simply because the model is so flexible. In other words, once a particular dataset is used to estimate the parameters of a model, the fit of the model on that sample is no longer an independent evaluation of the quality of the model. The most straightforward way to address this fit inflation would be testing the model on a new dataset. Another good way would be by repeated cross-validation (e.g., 5-fold, 10-fold or leave-one-out) using the existing dataset. However, AIC and similar criteria attempt to directly calculate an estimate of corrected fit (see Hastie *et al.*, 2009; Shao, 1993, 1997).

Akaike (1973) showed that an approximately unbiased estimate of *E_t_(l(y))* would be a constant plus 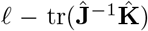 (where **J** and **K** are two *p × p* matrices, described below, and tr() is the trace, or sum of diagonal elements). 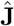 is an estimator for the covariance matrix of the parameters, based on the matrix of second derivatives of *ℓ* in each of the parameters, and 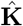 is an alternative estimator for the covariance matrix of the parameters, based on the cross-products of the first derivatives (see Claeskens and Hjort, 2008, pp. 26-7). Akaike showed that 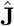 and 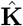 are asymptotically equal for the true model, so that the trace becomes approximately *p*, the number of parameters in the model. For models that are far from the truth, the approximation may not be as good. However, poor models presumably have poor values of *ℓ*, so the precise size of the penalty is less important (Burnham and Anderson, 2002). The resulting expression *ℓ* – *p* suggests using *A_n_* = 2 in (1) and concluding that fitted models with low values of (1) will be likely to provide a likelihood function closer to the truth.

#### 1.1.2 Criteria Related to AIC

When *n* is small or *p* is large, the crucial AIC approximation 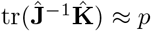 is too optimistic and the resulting penalty for model complexity is too weak (Tibshirani and Knight, 1999; Hastie *et al.*, 2009). In the context of linear regression and time series models, several researchers (e.g., Sugiura, 1978; Hurvich and Tsai, 1989; Burnham and Anderson, 2004) have suggested using a corrected version, AICc, which applies a slightly heavier penalty that depends on *p* and *n*; it gives results very close to those of AIC when *n/p* is large. The AIC_c_ can be written as 1 with *A_n_* = 2*n*/(*n* – *p* – 1). Theoretical discussions of model selection have often focused on asymptotic comparisons for large *n* and small *p*, and AICc gets little attention in this setting because it becomes equivalent to AIC as *n/p* → ∞. However, this equivalence is subject to the assumption that *p* is fixed and *n* becomes very large. Because in many situations *p* is comparable to *n* or larger, AICc may deserve more attention in future work.

Some other selection approaches are asymptotically equivalent for selection purposes to AIC, at least for linear regression. That is, they select the same model as AIC with high probability if *n/p* is very high. These include Mallows’ *C_p_* (see George, 2000), leave-one-out cross-validation (Shao, 1997; Stone, 1977), and the generalized cross-validation (GCV) statistic (see Golub *et al.*, 1979; Hastie *et al.*, 2009). Leave-one-out cross-validation involves fitting the candidate model on many subsamples of the data, each excluding one subject (i.e., participant or specimen), and observing the average squared error in predicting the extra response. Each approach is intended to correct a fit estimate for the artificial inflation in observed performance caused by fitting a model and evaluating it with the same data, and to find a good balance between bias caused by too restrictive a model and excessive variance caused by a model with too many parameters (Hastie *et al.*, 2009). These AIC-like criteria do not treat model parsimony as a motivating goal in its own right, but only as a means to reduce unnecessary sampling error caused by having to estimate too many parameters relative to *n*. Thus, especially for large *n*, AIC-like emphasize sensitivity more than specificity. However, in many research settings, parsimonious interpretation is of strong interest in its own right. In these settings, another criterion such as BIC, described in the next section, might be more appropriate.

Some other, more ad-hoc criteria are named after AIC but do not derive from the same theoretical framework, except that they share the form (1). For example, some researchers (Andrews and Currim, 2003; Fonseca and Cardoso, 2007; Yang and Yang, 2007) have suggested using *A_n_* = 3 in expression (1) instead of 2. The use of *A_n_* = 3 is sometimes called “AIC3.” There is no statistical theory to motivate AIC3, such as minimizing KL divergence or any other theoretical construct, but on an *ad hoc* basis it has fairly good simulation performance in some settings, being stricter than AIC but not as strict as BIC. Also, the CAIC, the “corrected AIC” or “consistent AIC” proposed by Bozdogan (1987), uses *A_n_* = ln(*n*) + 1. (It should not be confused with the AIC_c_ discussed above.) This penalty tends to result in a more parsimonious model and more underfitting than AIC or even than BIC. This value of *A_n_* was chosen somewhat arbitrarily as an example of an *A_n_* that would provide model selection consistency, a property described below in the section for BIC. However, any *A_n_* proportional to ln(*n*) provides model selection consistency, so CAIC has no real advantage over the better-known and better-studied BIC (see below), which also has this property.

Another of the “information criteria” commonly used in model selection, namely the Deviance Information Criterion (DIC) used in Bayesian analyses (Gibson *et al.*, 2018; Spiegelhalter *et al.*, 2002), cannot be expressed as a special case of Expression (1). It has a close relationship to AIC and has an analogous purpose within some Bayesian analyses (Ando, 2013; Claeskens and Hjort, 2008) but is conceptually and practically different and more complicated to compute. It is beyond the scope of this review because it is usually not used in the same settings as the AIC, BIC, and other common criteria, so it is usually not a direct competitor with them.

#### 1.1.3 Schwarz’s Bayesian Information Criterion (BIC)

In Bayesian model selection, a prior probability is set for each model *M_i_*, and prior distributions (often uninformative priors for simplicity) are also set for the nonzero coefficients in each model. If we assume that one and only one model, along with its associated priors, is true, we can use Bayes’ theorem to find the posterior probability of each model given the data. Let Pr(*M_i_*) be the prior probability set by the researcher, and let Pr(y|*M_i_*) be the probability density of the data given *M_i_*, calculated as the expected value of the likelihood function of y given the model and parameters, over the prior distribution of the parameters. According to Bayes’ theorem, the posterior probability Pr(M_i_|y) of a model is proportional to Pr(*M_i_*) Pr(y|*M_i_*). The degree to which the data support *M_i_* over another model *M_j_* is given by the ratio of the posterior odds to the prior odds:

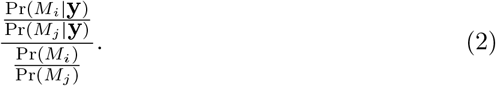

If we assume equal prior probabilities for each model, this simplifies to the “Bayes factor” (see Kass and Raftery, 1995):

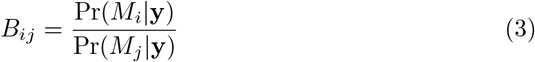

so that the model with the highest Bayes factor also has the higher posterior probability. Schwarz (1978) and Kass and Wasserman (1995) showed that, for many kinds of models, *B_ij_* can be roughly approximated by exp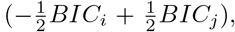, where BIC equals Expression (1) with *A_n_* = ln(*n*). BIC is also called the Schwarz criterion. Note that in a Bayesian analysis, all of the parameters within each of the candidate models have prior distributions representing knowledge or beliefs which the investigators have about their values before doing the study. The use of BIC assumes that a relatively noninformative prior is used, meaning that the prior is not allowed to have a large effect on the estimate of the coefficients (Kass and Wasserman, 1995; Weakliem, 1999). Thus, although Bayesian in origin, the BIC is often used in non-Bayesian analyses because it uses relatively noninformative priors which do not have to be set by the user. For fully Bayesian analyses with informative priors, posterior model probabilities or the previously mentioned Deviance Information Criterion (DIC) might be more appropriate.

The use of Bayes factors or their BIC approximation can be more interpretable than that of significance tests in some practical settings (Beard *et al.*, 2016; Goodman, 2008; Held and Ott, 2018; Raftery, 1996). BIC is described further in Raftery (1995) and Wasserman (2000), but critiqued by Gelman and Rubin (1995) and Weakliem (1999), who find it to be an oversimplification of Bayesian methods. Indeed, if Bayes factors or the BIC are used in an automatic way for choosing a single supposedly best model (e.g., setting a particular cutoff for choosing the larger model), then they are potentially subject to the same criticisms as classic significance tests (see Gigerenzer and Marewski, 2015; Murtaugh, 2014). However, Bayes factors or information criteria, if used thoughtfully, provide a way of comparing the appropriateness of each of a set of models on a common scale.

#### 1.1.4 Criteria Related to BIC

Sclove (1987) suggested a sample-size-adjusted BIC, variously abbreviated as ABIC, SABIC, or BIC*, based on the work of Rissanen (1978) and Boekee and Buss (1981). It uses *A_n_* = ln((*n*+2)/24) instead of *A_n_* = ln(*n*). This penalty will be much lighter than that of BIC, and may be lighter or heavier than that of AIC, depending on *n*. The unusual expression for *A_n_* comes from Rissanen’s work on model selection for autoregressive time series models from a minimum description length perspective (see Stine, 2004). It is not clear whether or not the same adjustment is still theoretically appropriate in different contexts, but in practice it is sometimes used in latent class modeling and seems to work fairly well (see Nylund *et al.*, 2007; Tein *et al.*, 2013). Table 2 gives the values *A_n_* for AIC, ABIC, BIC and CAIC for some representative values of *n*. It shows that CAIC always has the strongest penalty function. BIC has a stronger penalty than AIC for reasonable values of *n*. The ABIC has the property of usually being stricter than AIC but not as strict as BIC, which may be appealing to some researchers, but unfortunately it does not always really “adjust” for the sample size. In fact, for very small *n*, ABIC has a nonsensical negative penalty encouraging needless complexity. AICc is not shown in the table because its *A_n_* depends on *p* as well as *n*.

**Table 2:** *A_n_* for Common Information Criteria

| $n$ | AIC | ABIC | BIC | CAIC |
| --- | --- | --- | --- | --- |
| 10 | 2.0000 | -0.6931 | 2.3026 | 3.3026 |
| 50 | 2.0000 | 0.7732 | 3.9120 | 4.9120 |
| 100 | 2.0000 | 1.4469 | 4.6052 | 5.6052 |
| 500 | 2.0000 | 3.0405 | 6.2146 | 7.2146 |
| 1000 | 2.0000 | 3.7317 | 6.9078 | 7.9078 |
| 5000 | 2.0000 | 5.3395 | 8.5172 | 9.5172 |
| 10000 | 2.0000 | 6.0325 | 9.2103 | 10.2103 |
| 100000 | 2.0000 | 8.3349 | 11.5129 | 12.5129 |

**Notes.** $A_n$ = penalty weighting constant. $n$ = sample size (number of subjects). AIC = Akaike information criterion. ABIC = adjusted Bayesian information criterion. BIC = Bayesian information criterion. CAIC = Consistent Akaike information criterion.

### 1.2 AIC versus BIC and the Concept of Consistent Model Selection

BIC is sometimes preferred over AIC because BIC is “consistent” (e.g., Nylund *et al.*, 2007). Assuming that a fixed number of models are available and that one of them is the true model, a consistent selector is one that selects the true model with probability approaching 100% as *n* → ∞ (see Rao and Wu, 1989; Zhang, 1993; Shao, 1997; Yang, 2005; Claeskens and Hjort, 2008).

The existence of a true model here is not as unrealistically dogmatic as it sounds (Burnham and Anderson, 2004; Kuha, 2004). Rather, the *true* model can be defined as the simplest adequate model, that is, the single model that minimizes KL divergence, or the one such model with the fewest parameters if there is more than one (Claeskens and Hjort, 2008). There may be more than one such model because if a given model has a given KL divergence from the truth, any more general model containing it will have no greater distance from the truth. This is because there is some set of parameters for which the larger model becomes the model nested within it. However, the theoretical properties of BIC are better in situations in which a model with a finite number of parameters can be treated as “true” (Shao, 1997). In summary, even though at first the BIC seems fraught with philosophical problems because of its apparent assumption of that one of the models available is the “true” one, it is nonetheless well-defined and useful in practice.

AIC is not consistent because it has a non-vanishing chance of choosing an unnecessarily complex model as *n* becomes large. The unnecessarily complex model would still closely approximate the true distribution but would use more parameters than necessary to do so. However, selection consistency involves some performance tradeoffs when *n* is modest, specifically, an elevated risk of poor performance caused by underfitting (see Potscher, 1991; Shao, 1997; Shibata, 1986; Vrieze, 2012). In general, the strengths of AIC and BIC cannot be combined by any single choice of *A_n_* (Leeb, 2008; Yang, 2005). However, in some cases it is possible to construct a more complicated model selection approach that uses aspects of both (see Ding *et al.*, 2018).

Nylund *et al.* (2007) seem to interpret the lack of selection consistency as a flaw in AIC (Nylund *et al.*, 2007, p. 556). However, we argue the real situation is somewhat more complicated; AIC is not a defective BIC, nor *vice versa* (see Potscher, 1991; Vrieze, 2012). Likewise, the other ICs mentioned here are neither right nor wrong, but are simply choices (perhaps thoughtful and perhaps arbitrary, but still technically valid choices).

## 2 Information Criteria in Simple Cases

AIC and BIC differ in theoretical basis and interpretation (Aho *et al.*, 2014; Claeskens and Hjort, 2008; Kuha, 2004; Shmueli, 2010). They also sometimes disagree in practice, generally with AIC indicating models with more parameters and BIC with less. This has led many researchers to question whether and when a particular value of the “magic number” *A_n_* (Bozdogan, 1987) can be chosen as most appropriate. Two special cases – comparing equally sized models and comparing nested models – each provide some insight into this question.

First, *when comparing different models of the same size* (i.e., number of parameters to be estimated), all ICs of the form (1) will always agree on which model is best. For example, in regression variable subset selection, suppose two models each use five covariates. In this case, any IC will select whichever model has the highest likelihood (the best fit to the observed sample) after estimating the parameters. This is because only the first term in Expression (1) will differ across the candidate models, so *A_n_* does not matter. Thus, although the ICs differ in theoretical framework, they only disagree when they make different tradeoffs between fit and model size.

Second, *for comparing a nested pair of models, different ICs act like different α levels on a likelihood ratio test* (LRT). For comparing models of different sizes, when one model is a restricted case of the other, the larger model will typically offer better fit to the observed data at the cost of needing to estimate more parameters. The ICs will differ only in how they make this bias-variance tradeoff (Lin and Dayton, 1997; Sclove, 1987). Thus, an IC will act like a hypothesis test with a particular α level (Claeskens and Hjort, 2008; Derryberry *et al.*, 2018; Foster and George, 1994; Murtaugh, 2014; Potscher, 1991; Söderström, 1977; Stoica *et al.*, 2004; van der Hoeven, 2005; Vrieze, 2012).

Suppose a researcher will choose whichever of *M_0_* and *M_1_* has the better (lower) value of an IC of the form (1). This means that *M_1_* will be chosen if and only if –2*ℓ_1_+A_n_p*_1_ < –2*ℓ*_0_+*A_n_p*_0_, where *ℓ*_1_ and *ℓ*_0_ are the fitted maximized log-likelihoods for each model. Although the comparison of models is interpreted differently in the theoretical frameworks used to justify AIC and BIC (Aho *et al.*, 2014; Kuha, 2004), algebraically this comparison is the same as a LRT (Potscher, 1991; Söderström, 1977; Stoica *et al.*, 2004). That is, *M*_0_ is rejected if and only if

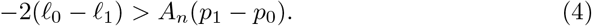

The left-hand side is the LRT test statistic (since a logarithm of a ratio of quantities is the difference in the logarithms of the quantities). Thus, in the case of nested models an IC comparison is mathematically an LRT with a different interpretation. The *α* level is specified indirectly through the critical value *A_n_*; it is the proportion of the null hypothesis distribution of the LRT statistic that is less than *A_n_*.

### 2.1 Implications of the LRT Equivalence in the Nested Case

For comparing nested maximum-likelihood models satisfying classic regularity conditions, including classical linear and logistic regression models (although not necessarily including mixture models; see Chernoff and Lander, 1995; McLachlan and Peel, 2000) the null-hypothesis distribution of –2(*ℓ*_0_ – *ℓ*_1_) is asymptotically χ^2^ with degrees of freedom (*df*) equal to *p*_1_ – *p*_0_. Consulting a χ^2^ distribution and assuming *p*_1_ – *p*_0_ = 1, AIC (*A_n_* = 2) becomes equivalent to a LRT test at an *α* level of about .16 (i.e., the probability of a 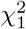 deviate being greater than 2). For example, in the case of linear regression, comparing IC’s of otherwise identical models differing by the presence or absence of a covariate can also be shown to be mathematically equivalent to a significance test for the regression coefficient of that covariate (Derryberry *et al.*, 2018).

In the same situation, BIC (with *A_n_* = ln(*n*)) has an *α* level that depends on *n*. If *n* = 10 then *A_n_* = ln(*n*) = 2.30 so *α* = .13. If *n* = 100 then *A_n_* = 4.60 so *α* = .032. If *n* = 1000 then *A_n_* = 6.91 so *α* = .0086, and so on. Thus when *p*_1_ – *p*_0_ = 1, significance testing at the customary level of *α* = .05 is often an intermediate choice between AIC and BIC, corresponding to *A_n_* = 1.96^2^ ∈ 4. However, as *p*_1_ – *p*_0_ becomes larger, all ICs become more conservative, in order to avoid adding many unnecessary parameters unless they are needed. Table 3 shows different effective α values for two values of *p*_1_ – *p*_0_, obtained using the R (R Core Team, 2017) code `1-pchisq(q=An*df,df=df,lower.tail=TRUE)` where `An` is the *A_n_* value and `df` is *p*_1_ – *p_0_.* AIC_c_ is not shown in the table because its penalty weight depends both on *p*_0_ and on *p*_1_ in a slightly more complicated way, but will behave similarly to AIC for large n and modest *p*_0_.

**Table 3:** Alpha Levels Represented By Common Information Criteria

| $n$ | AIC | ABIC | BIC | CAIC |
| --- | --- | --- | --- | --- |
| Assuming $p_1 - p_0 = 1$ | | | | |
| 10 | 0.15730 | 1.00000 | 0.12916 | 0.06917 |
| 50 | 0.15730 | 0.37923 | 0.04794 | 0.02667 |
| 100 | 0.15730 | 0.22902 | 0.03188 | 0.01791 |
| 500 | 0.15730 | 0.08121 | 0.01267 | 0.00723 |
| 1000 | 0.15730 | 0.05339 | 0.00858 | 0.00492 |
| 5000 | 0.15730 | 0.02085 | 0.00352 | 0.00204 |
| 10000 | 0.15730 | 0.01404 | 0.00241 | 0.00140 |
| 100000 | 0.15730 | 0.00389 | 0.00069 | 0.00040 |
| Assuming $p_1 - p_0 = 10$ | | | | |
| 10 | 0.02925 | 1.00000 | 0.01065 | 0.00027 |
| 50 | 0.02925 | 0.65501 | 0.00002 | < 0.00001 |
| 100 | 0.02925 | 0.15265 | < 0.00001 | < 0.00001 |
| 500 | 0.02925 | 0.00074 | < 0.00001 | < 0.00001 |
| 1000 | 0.02925 | 0.00005 | < 0.00001 | < 0.00001 |
| 5000 | 0.02925 | < 0.00001 | < 0.00001 | < 0.00001 |
| 10000 | 0.02925 | < 0.00001 | < 0.00001 | < 0.00001 |
| 100000 | 0.02925 | < 0.00001 | < 0.00001 | < 0.00001 |

**Notes.** $n$ = sample size (number of subjects). AIC = Akaike information criterion. ABIC = adjusted Bayesian information criterion. BIC = Bayesian information criterion. CAIC = Consistent Akaike information criterion. $p_1$ = number of free parameters in larger model within pair being compared. $p_0$ = number of free parameters in smaller model.

### 2.2 Interpretation of Selection Consistency

The property of selection consistency can be intuitively understood from this perspective. For AIC, as for hypothesis tests, the power of a test typically increases with *n* because *ℓ*_1_ and *ℓ*_0_ are sums over the entire sample. This is why empirical studies are planned to have adequate sample size to guarantee a reasonable chance of success (Cohen *et al.*, 2003). Unfortunately rejecting any given false null hy-pothesis is practically guaranteed for sufficiently large *n* even if the effect size is tiny. However, the Type I error rate is constant and never approaches zero. On the other hand, BIC becomes a more stringent test (has a decreasing Type I error rate) as *n* increases. The power increases more slowly (i.e., the Type II error rate decreases more slowly) than for AIC or for fixed-*α* hypothesis tests because the test is becoming more stringent, but now the Type I error rate is also decreasing. Thus, nonzero but practically negligible departures from a model are less likely to lead to rejecting the model for BIC than for AIC (Raftery, 1995). Fortunately, even for BIC, the decrease in *α* as *n* increases is slow; thus power still increases as *n* increases, although more slowly than it would for AIC. Thus, for BIC, both the Type I and Type II error rates decline slowly as *n* increases, while for AIC (and for classical significance testing) the Type II error rate declines more quickly but the Type I error rate does not decline at all. This is intuitively why a criterion with constant *A_n_* cannot be asymptotically consistent even though it may be more powerful for a given *n* (see Claeskens and Hjort, 2008; Yang, 2005; Derryberry *et al.*, 2018).

Also, since choosing *A_n_* for a model comparison is closely related to choosing an *α* level for a significance test, it becomes clear that the universally “best” IC cannot be defined any more than the “best” *α*; there will always be a tradeoff. Thus, debates about whether AIC is generally superior to BIC or *vice versa*, will be fruitless.

### 2.3 Interpretation in Terms of Tradeoffs

*For non-nested models of different sizes*, neither of the above simple cases hold; furthermore, these complex cases are often those in which ICs are most important because a LRT cannot be performed. However, it remains the case that *A_n_* indirectly controls the tradeoff between the likelihood term and the penalty on the number of parameters, hence the tradeoff between good fit to the observed data and parsimony.

Almost by definition, there is no universal best way to decide how to make a tradeoff. Type I errors are generally considered worse than Type II errors, because the former involve introducing false findings into the literature while the latter are simply non-findings. However, Type II errors involve the loss of potentially important scientific discoveries, and furthermore both kinds of errors can lead to poor policy or treatment decisions in practice, especially because failure to reject *H*_0_ is often misinterpreted as demonstrating the truth of *H*_0_ (Peterman, 1990). Thus, researchers try to specify a reasonable *α* level which is neither too low (causing low power) nor too high (inviting false positive findings). In this way, model comparison is much like a medical diagnostic test (see, e.g., Altman and Bland, 1994), replacing “Type I error” with “false positive” and “Type II error” with “false negative.” AIC and BIC use the same data but apply different cutoffs for whether to “diagnose” the smaller model as being inadequate. AIC is more sensitive (lower false-negative rate), but BIC is more specific (lower false-positive rate). The utility of each cutoff is determined by the consequences of a false positive or false negative and by one’s beliefs about the base rates of positives and negatives. Thus, AIC and BIC could be seen as representing different sets of prior beliefs in a Bayesian sense (see Burnham and Anderson, 2004; Kadane and Lazar, 2004) or, at least, different judgments about the importance of parsimony. Perhaps in some examples a more or less sensitive test (higher or lower *A_n_* or *α*) would be more appropriate than in others. For example, although AIC has favorable theoretical properties for choosing the number of parameters needed to approximate the shape of a nonparametric growth curve in general (Shao, 1997), in a particular application with such data Dziak *et al.* (2015) argued that BIC would give more interpretable results. They argued this because the curves in that context were believed likely to have a smooth and simple shape, as they represented averages of trajectories of an intensively measured variable on many individuals with diverse individual experiences and because deviations from the trajectory could be modeled using other aspects of the model.

However, in practice it is often difficult to determine the *α* value that a particular criterion really represents, for two reasons. First, even for regular situations in which a LRT is known to work well, the χ^2^ distribution for the test statistic is asymptotic and will not apply well to small *n*. Second, in some situations the rationale for using an IC is, ironically, the failure of the assumptions needed for a LRT. That is, the test emulated by the IC will itself not be valid at its nominal *α* level anyway. Therefore, although the comparison of *A_n_* to an *α* level is helpful for getting a sense of the similarities and differences among the ICs, simulations are required to describe exactly how they behave. In the section below we review simulation results from a common application of ICs, namely the selection of the number of latent classes (empirically derived clusters) in a dataset.

## 3 The Special Case of Latent Class Analysis

A common use of ICs is in selecting the number of components for a latent class analysis (LCA). LCA is a kind of finite mixture model (essentially, a model-based cluster analysis; McLachlan and Peel, 2000; Lazarsfeld and Henry, 1968; Collins and Lanza, 2010). LCA assumes that the population is a “mixture” of multiple classes of a categorical latent variable. Each class has different parameters that define the distributions of observed items, and the goal is to account for the relationships among items by defining classes appropriately. LCA is very similar to cluster analysis, but is based on maximizing an explicitly stated likelihood function rather than focusing on a heuristic computational algorithm like k-means. Also, some authors use the term LCA only when the observed variables are also categorical (as in the cancer symptoms example described above), and use the term “latent profile analysis” for numerical observed variables (as in the iris example), but we ignore this distinction here. LCA is also closely related to latent transition (LTA) models (see Collins and Lanza, 2010), an application of hidden Markov models (see, e.g., Eddy, 2004) that allows changes in latent class membership, conceptualized as transitions in an unobserved Markov chain. LCA models are sometimes used in combination with other models, such as in predicting class membership from genotypic or demographic variables, or predicting medical or behavioral phenotypes from class membership (e.g., Bray *et al.*, 2018; Dziak *et al.*, 2016; Lubke *et al.*, 2012).

For a simple latent class analysis without additional covariates, there are two kinds of model parameters: the sizes of the classes, and the class-specific parameters. For binary outcomes as in the cancer symptoms study, there is a class-specific parameter for each combination of class and item, giving the probability of endorsing this item given membership in this class. For numerical outcomes, the means and covariance parameters of the vector of items within each class constitute the class-specific parameters. To fit an LCA model or any of its cousins, an algorithm such as EM (Dempster *et al.*, 1977; Gupta and Chen, 2010; McLachlan and Peel, 2000) is often used to alternatively estimate class-specific parameters and predict subjects’ class membership given those parameters. The user must specify the number of classes in a model, but the true number of classes is generally unknown (Nylund *et al.*, 2007; Tein *et al.*, 2013). Sometimes one might have a strong theoretical reason to specify the number of classes, but often this must be done using data-driven model selection.

### 3.1 ICs for Selecting the Number of Classes

A naïve approach would be to use likelihood ratio (LR) or deviance (*G^2^*) tests sequentially to choose the number of classes and to conclude that the *k*-class model is large enough if and only if the (*k*+1)-class model does not fit the data significantly better. The selected number of classes would be the smallest *k* that is not rejected when compared to the (*k* + 1)-class model. However, the assumptions for the supposed asymptotic χ^2^ distribution in a LRT are not met in the setting of LCA, so that the *p*-values from those tests are not valid (see Lin and Dayton, 1997; McLachlan and Peel, 2000). The reasons for this are based on the fact that here is not nested in a regular way within H_1_, since a *k*-class model is obtained from a (*k* + 1)-class model either by constraining any one of the class sizes to a boundary value of zero or by setting the class-specific item-response probabilities equal between any two classes. That is, an meaningful *k*-class model is not obtained simply by setting a parameter to zero in a (*k* +1) class model in the way that, for example, a more parsimonious regression model can be obtained by starting with a model with many covariates and then constraining certain coefficients to zero. Ironically, the lack of regular nesting structure that makes it impossible to decide on the number of classes with an LRT has also been shown to invalidate the mathematical approximations used in the AIC and BIC derivations in the same way (McLachlan and Peel, 2000, pp. 202-212). Nonetheless, ICs are widely used in LCA and other mixture models. This is partly due to their ease of use, even without a firm theoretical basis. Fortunately, there is at least an asymptotic theoretical result showing that, when the true model is well-identified, BIC (and hence also AIC and ABIC) will have a probability of underestimating the true number of classes that approaches 0 as sample size tends to infinity (Leroux, 1992; McLachlan and Peel, 2000, p. 209).

### 3.2 Past Simulation Studies

Lin and Dayton (1997) did an early simulation study comparing the performance of AIC, BIC, and CAIC for choosing which assumptions to make in constructing constrained LCA models, a model selection task which is somewhat but not fully analogous to choosing the number of classes. When a very simple model was used as the true model, BIC and CAIC were more likely to choose the true model than AIC, which tended to choose an unnecessarily complicated one. When a more complex model was used to generate the data and measurement quality was poor, AIC was more likely to choose the true model than BIC or CAIC, which were likely to choose an overly simplistic one. They explained that this was very intuitive given the differing degrees of emphasis on parsimony. Interpreting these results, Dayton (1998) suggested that AIC tended to be a better choice in LCA than BIC, but recommended computing and comparing both.

Other simulations have explored the ability of the ICs to determine the correct number of classes. In Dias (2006), AIC had the lowest rate of underfitting but often overfit, while BIC and CAIC practically never overfit but often underfit. AIC3 was in between and did well in general. The danger of underfitting increased when the classes did not have very different response profiles and were therefore easy to mistakenly lump together; in these cases BIC and CAIC almost always underfit. Yang (2006) reported that ABIC performed better in general than AIC (whose model selection accuracy never got to 100% regardless of *n*) or BIC or CAIC (which underfit too often and required large *n* to be accurate). Fonseca and Cardoso (2007) similarly suggested AIC3 as the preferred selection criterion for categorical LCA models.

Yang and Yang (2007) compared AIC, BIC, AIC3, ABIC and CAIC. When the true number of classes was large and *n* was small, CAIC and BIC seriously underfit, but AIC3 and ABIC performed better. Nylund *et al.* (2007) presented various simulations on the performance of various ICs and tests for selecting the number of classes in LCA, as well as factor mixture models and growth mixture models. Overall, in their simulations, BIC performed much better than AIC, which tended to overfit, or CAIC, which tended to underfit (Nylund *et al.*, 2007, p. 559). However, this does not mean that BIC was the best in every situation. In most of the scenarios considered by Nylund *et al.* (2007), BIC and CAIC almost always selected the correct model size, while AIC had a much smaller accuracy in these scenarios because of a tendency to overfit. In those scenarios, *n* was large enough so that the lower sensitivity of BIC was not a problem. However, in a more challenging scenario with a small sample and unequally sized classes, (Nylund *et al.*, 2007, p. 557), BIC essentially never chose the larger correct model and it usually chose one that was much too small. Thus, as Lin and Dayton (1997) found, BIC may select too few classes when the true population structure is complex but subtle (for example, a small but nonzero difference between the parameters of a pair of classes) and *n* is small. Wu (2009) compared the performance of AIC, BIC, ABIC, CAIC, naïve tests, and the bootstrap LRT in hundreds of simulated scenarios. Performance was heavily dependent on the scenario, but the method that worked adequately in the greatest variety of situations was the bootstrap LRT, followed by ABIC and classic BIC. Wu (2009) argued that BIC seemed to outperform ABIC in the most optimal situations because of its parsimony, but that ABIC seemed to do better in situations with smaller *n* or more unequal class sizes. Dziak *et al.* (2014) also concluded that BIC could seriously underfit relative to AIC for small sample sizes or other challenging situations. In latent profile analysis, Tein *et al.* (2013) found that BIC and ABIC did well for large sample sizes and easily distinguishable classes, but AIC chose too many classes, and no method performed well for especially challenging scenarios. In a more distantly related mixture modeling framework involving modeling evolutionary rates at different genomic sites, Kalyaanamoorthy *et al.* (2017) found that AIC, AIC_c_, and BIC worked well but that BIC worked best.

### 3.3 Difficulties of Applying Simulation Results

Despite all these findings, is not possible to say which IC is universally best, even in the idealized world of simulations. What constitutes a “large” or “small” n, for the purposes of the performance of BIC, depends on the true class sizes and characteristics, which by definition are unknown. For example, if there are many small classes, a larger overall sample size is needed to distinguish them all. A smaller number of flowers might have been needed in our flower example if there had been three genera instead of three species, and a larger number might be needed to distinguish three cultivars or subspecies. Thus, the point at which the *n* becomes “large” depends on numerous aspects of the simulated scenario (Brewer *et al.*, 2016; Dziak *et al.*, 2014). Furthermore, in real data, unlike simulations, the characteristics of the “true” (data-generating) model are unknown, since the data have been generated by a natural or experimental process rather than a probability model. For this reason it may be more helpful to think about which aspects of performance (e.g., sensitivity or specificity) are most important in a given situation, rather than talking about the nature of a supposed true data-generating model.

If the goal of having a model which contains enough parameters to describe the heterogeneity in the population is more important than the goal of parsimony, or if some classes are expected to be small or similar to other classes but distinguishing among them is still considered important for theoretical reasons, then perhaps AIC, AIC3 or ABIC should be used instead of BIC or CAIC. If obtaining a few large and distinctly interpretable classes is more important, then BIC is more appropriate. Sometimes, the AIC-favored model might be so large as to be difficult to use or understand. In these cases, the BIC-favored model is clearly the better practical choice. For example, in Chan *et al.* (2007) BIC favored a mixture model with 5 classes, and AIC favored at least 10; the authors felt that a 10-class model would be too hard to interpret. In fact, it may be necessary for theoretical or practical reasons to choose a number of classes even smaller than that suggested by BIC. This is because it is important to choose the number of classes based on their theoretical interpretability, as well as by excluding any models with so many classes that they lead to a failure to converge to a clear maximum-likelihood solution (see Bray and Dziak, 2018; Collins and Lanza, 2010; Pohle *et al.*, 2017).

### 3.4 Other Methods for Selecting the Number of Classes

An alternative to ICs in latent class analysis and cluster analysis is the use of a bootstrap test (see McLachlan and Peel, 2000). Unlike the naïve NRT, Nylund *et al.* (2007) showed empirically that the bootstrap LRT with a given *α* level does generally provide a Type I error rate at or below that specified level. Both Nylund *et al.* (2007) and Wu (2009) found that this bootstrap test seemed to perform somewhat better than the ICs in various situations. The bootstrap LRT is beyond the scope of this paper, as are more computationally intensive versions of AIC and BIC, involving bootstrapping, cross-validation, or posterior simulation (see McLachlan and Peel, 2000, pp. 204-212). Also beyond the scope of this paper are mixture-specific selection criteria such as the normalized entropy criterion (Biernacki *et al.*, 1999) or integrated completed likelihood (Biernacki and Celeux, 2000; Rau and Maugis, 2018), or the minimum message length approach of Silvestre *et al.* (2014). However, the basic ideas in this article will still be helpful in interpreting the implications of some of the other selection methods. For example, like any test or criterion, the bootstrap LRT still requires the choice of a tradeoff between sensitivity and specificity (i.e., by selecting an *α* level).

## 4 Discussion

Many simulation studies have been performed to compare the performance of information criteria. For small *n* or difficult-to-distinguish classes, the most likely error in a simulation is underfitting, so the criteria with lower underfitting rates, such as AIC, often seem better. For very large *n* and easily distinguished classes, the most likely error is overfitting, so more parsimonious criteria, such as BIC, often seem better. However, the true model structure, parameter values, and sample size used when generating simulated data determine the relative performance of the ICs in simulations in a complicated way, limiting the extent to which they can be used to state general rules or advice (Brewer *et al.*, 2016; Dziak *et al.*, 2014; Emiliano *et al.*, 2014).

If BIC indicates that a model is too small, it may well be too small (or else fit poorly for some other reason). If AIC indicates that a model is too large, it may well be too large for the data to warrant. Beyond this, theory and judgment are needed. If BIC selects the largest and most general model considered, it is worth thinking about whether to expand the space of models considered (since an even more general model might fit even better), and similarly if AIC chooses the most parsimonious.

AIC and BIC each have distinct theoretical advantages. However, a researcher may judge that there may be a practical advantage to one or the other in some situations. For example, as mentioned earlier, in choosing the number of classes in a mixture model, the true number of classes required to satisfy all model assumptions is sometimes quite large, too large to be of practical use or even to allow coefficients to be reliably estimated. In that case, BIC would be a better choice than AIC. Additionally, in practice, one may wish to rely on substantive theory or parsimony of interpretation in choosing a relatively simple model. In such cases, the researcher may decide that even the BIC may have indicated a model that is too complex in a practical sense, and may choose to select a smaller model that is more theoretically meaningful or practically interpretable instead (Bray and Dziak, 2018; Pohle *et al.*, 2017). This does not mean that BIC overfit. Rather, in these situations the model desired is sometimes not the literally true model but simply the most useful model, a concept which cannot be identified using fit statistics alone but requires subjective judgment. Depending on the situation, the number of classes in a mixture model may either be interpreted a true quantity needing to be objectively estimated, or else as a level of approximation to be chosen for convenience, like the scale of a map. Still, in either case the question of which patterns or features are generalizable beyond the given sample remains relevant (c.f. Li and Marron, 2005). In the iris example, there was a consensus correct answer given by the number of recognized biological species. However, in the cancer symptoms example, the latent classes were more a convenient way of summarizing the data than a reflection of distinct underlying syndromes. If a fifth class had been included, it might have been something like “moderate physical, moderate psychological” which probably would not have provided additional insights beyond those which could be gained by comparing the four classes in the four-class model. Of course, in some studies, classes or trajectories might represent different biological processes of distinct clinical importance (e.g., Karlsson *et al.*, 2018), and then it might be very important not to miss any, but in other cases they may simply be regions in an underlying multivariate continuum.

One could use the ICs to suggest a range of model sizes to consider for future study; for example, in some cases one might use the BIC-preferred model as a minimum size and the AIC-preferred model as a maximum. Either AIC or BIC can also be used for model averaging, that is, estimating quantities of interest by combining more than one model weighted by their plausibility (see Burnham and Anderson, 2004; Claeskens and Hjort, 2008; Gelman and Rubin, 1995; Hoeting *et al.*, 1999; Johnson and Omland, 2004; Minin *et al.*, 2003; Posada and Crandall, 2001; Posada and Buckley, 2004).

Although model selection is not an entirely objective process, it can still be a scientific one (see Gelman and Hennig, 2017). The fact that there is no universal consensus on a way to choose a model is not a bad thing; an automatic and uncritical use of an IC is no more insightful than an automatic and uncritical use of a *p*-value (Brewer *et al.*, 2016; Emiliano *et al.*, 2014; Gigerenzer and Marewski, 2015). Comparing different information criteria may suggest what range of models is reasonable. Of course, researchers must explain their methodological choices and not pick and choose methods simply as a way of supporting a desired outcome (see Simmons *et al.*, 2011).

A larger question is whether to use ICs at all. If ICs indeed reduce to LRTs in simple cases, one might wonder why ICs are needed at all, and why researchers cannot simply do LRTs. A possible answer is flexibility. Both AIC and BIC can be used to concurrently compare many models, not all of them nested, rather than just a pair of nested models at a time. They can also be used to weight the estimates obtained from different models for a common quantity of interest. These weighting approaches use either AIC or BIC but not both, because AIC and BIC are essentially treated as different Bayesian priors. While currently we know of no mathematical theoretical framework for explicitly combining both AIC and BIC into a single weighting scheme, a sensitivity analysis could be performed by comparing the results from both. AIC and BIC can also be used to choose a few well-fitting models, rather than selecting a single model from among many and assuming it to be the truth (Kuha, 2004). Researchers have also proposed benchmarks for judging whether the size of a difference in AIC or BIC between models is practically significant (see Burnham and Anderson, 2004; Murtaugh, 2014; Raftery, 1995); for example, an AIC or BIC difference between two models of less than 2 provides little evidence for one over the other; an AIC or BIC difference of 10 or more is strong evidence. These principles should not be used as rigid cutoffs (Murtaugh, 2014), but as input to decision making and interpretation. Kadane and Lazar (2004) suggested that ICs might be used to “deselect” very poor models (p. 279), leaving a few good ones for further study, rather than indicating a single best model.

Consider a regression context in which we are considering variables A, B, C, D, and E; suppose also that the subset with the lowest BIC is {A,B,C} with a BIC of 34.2, while the second-best is {B,C,D} with a BIC of 34.3. A naïve approach would be to conclude that A is an important predictor and D is not, and then conduct all later estimates and analyses using only the subset { A,B,C}. If we had gathered an even slightly different sample, though, we might be just as likely to make the opposite conclusion. What should we do? Some researchers might just report one model as being the correct one and ignore the other. However, this seriously understates the true degree of uncertainty present (Burnham and Anderson, 2002). Considering more than one IC, such as AIC and BIC together, could make even more models seem plausible. A simple sequential testing approach with a fixed *α* =.05 would seemingly avoid this ambiguity. However, the avoidance of ambiguity there would be artificial – the uncertainty still exists but is being ignored.

In many cases, cross-validation approaches can be used as good alternatives to IC’s. However, they are sometimes more computationally intensive. Also, implementation details of the cross-validation approaches can affect parsimony in an analogous way to the choice of *A_n_* (Yang, 2007).

Lastly, both AIC and BIC were developed in situations in which *n* was assumed to be much larger than *p*. None of the ICs discussed here were specifically developed for situations such as those found in many genome-wide association studies predicting disease outcomes, in which the number of participants (*n*) is often smaller than the number of potential genes (*p*), even when *n* is in the tens of thousands. The ICs can still be practically useful in this setting (e.g., Cross-Disorder Group of the Psychiatric Genomics Consortium, 2013). However, sometimes they might need to be adapted (see, e.g., Chen and Chen, 2008; Liao *et al.*, 2018; Mestres *et al.*, 2018; Pan *et al.*, 2016). More research in this area would be worthwhile.

## Code Appendix

The R code below performs the cluster analysis and model selection described above for the iris data.

~~~
library(mclust);
library(datasets);
n <-150;
ll <-rep(NA,7);
bic.given <-rep(NA,7);
models <-list();
for (k in 1:7) {
   temp.model <-Mclust(iris[,1:4],G=k,modelNames=“VVV”);
   p[k] <-temp.model$df;
   ll[k] <-temp.model$loglik;
   bic.given[k] <-temp.model$bic;
   models[[k]] <-temp.model;
}
aic.calculated <- -2*ll + 2*p;
caic.calculated <- -2*ll + (1+log(n))*p;
abic.calculated <- -2*ll + log((n+2)/24)*p;
bic.calculated <- -2*ll + log(n)*p;
print(cbind(aic.calculated,bic.calculated,abic.calculated,caic.calculated));
table(predict(models[[2]])$classification,iris$Species)
table(predict(models[[3]])$classification,iris$Species)
~~~

## Acknowledgements

The authors thank Dr. Linda M. Collins for very valuable suggestions and insights which helped in the development of this paper. We also thank Dr. Michael Cleveland for his careful review and recommendations on an earlier version of this paper and Amanda Applegate for her suggestions. John Dziak thanks Frank Sabach for encouragement in the early stages of his research. Lars Jermiin thanks the University College Dublin for its generous hospitality.

A previous version of this report has been disseminated as Methodology Center Technical Report 12-119, June 27, 2012, and as a preprint at https://peerj.com/preprints/1103/. The earlier version of the paper contains simulations to illustrate the points made. Simulation code is available at http://www.runmycode.org/companion/view/1306 and results at https://methodology.psu.edu/media/techreports/12-119.pdf.

## Funding

This research was supported by NIH grant P50 DA039838 from the National Institute on Drug Abuse. The content is solely the responsibility of the authors and does not necessarily represent the official views of the National Institute on Drug Abuse or the National Institutes of Health.

